# A Step Towards Generalisability: Training a Machine Learning Scoring Function for Structure-Based Virtual Screening

**DOI:** 10.1101/2022.10.28.511712

**Authors:** Jack Scantlebury, Lucy Vost, Anna Carbery, Thomas E. Hadfield, Oliver M. Turnbull, Nathan Brown, Vijil Chenthamarakshan, Payel Das, Harold Grosjean, Frank von Delft, Charlotte M. Deane

## Abstract

Over the last few years, many machine learning-based scoring functions for predicting the binding of small molecules to proteins have been developed. Their objective is to approximate the distribution which takes two molecules as input and outputs the energy of their interaction. Only a scoring function that accounts for the interatomic interactions involved in binding can accurately predict binding affinity on unseen molecules. However, many scoring functions make predictions based on dataset biases rather than an understanding of the physics of binding. These scoring functions perform well when tested on similar targets to those in the training set, but fail to generalise to dissimilar targets. To test what a machine learning-based scoring function has learnt, input attribution—a technique for learning which features are important to a model when making a prediction on a particular data point—can be applied. If a model successfully learns something beyond dataset biases, attribution should give insight into the important binding interactions that are taking place. We built a machine learning-based scoring function that aimed to avoid the influence of bias via thorough train and test dataset filtering, and show that it achieves comparable performance on the CASF-2016 benchmark to other leading methods. We then use the CASF-2016 test set to perform attribution, and find that the bonds identified as important by PointVS, unlike those extracted from other scoring functions, have a high correlation with those found by a distance-based interaction profiler. We then show that attribution can be used to extract important binding pharmacophores from a given protein target when supplied with a number of bound structures. We use this information to perform fragment elaboration, and see improvements in docking scores compared to using structural information from a traditional, data-based approach. This not only provides definitive proof that the scoring function has learnt to identify some important binding interactions, but also constitutes the first deep learning-based method for extracting structural information from a target for molecule design.

## 1. Introduction

The recent explosion of machine learning (ML) across all scientific disciplines has been accompanied by concerns regarding the generalisability of methods used. Various studies have found data leakage to be present in ML applications, resulting in overly optimistic performances being reported for the tools in question.^1–4^ The field of drug discovery is by no means exempt from this: models that learn unintended features from training datasets are extremely sensitive to small dataset distribution shifts, and cannot make reliable predictions on out-of-distribution datapoints. Practically, this renders them incapable of assisting in the development of drugs for novel targets.^5–7^

Typically led by human experts, drug development is a time-consuming and expensive process, with recent estimates placing the time required to reach clinical trials at 8.3 years, and the median cost at $985 million.^8^ Computational techniques offer a promising alternate route to human-led design. One such computational technique is docking.^9,10^ Docking algorithms take as input the coordinates of protein and ligand atoms, and predict the possible conformations of the protein-ligand complex. There exists a function describing the energy of the complex, and as conformational space is searched, this function is minimised. In the case of a deterministic algorithm with a single starting conformation, a single output pose is generated along with an estimation of the binding affinity; for a probabilistic search, or any search with multiple starting conformations, the result is a list of binding poses. These are ranked in the order of estimated binding affinity. Affinities are calculated using a scoring function, which uses a combination of atomic interactions to approximate the binding energy.

Over the last decade, machine learning models have shown promise in predicting both the pose and energetics of protein-ligand binding. In particular, deep learning models, which can be made to approximate extremely complex functions,^11^ have been used to predict binding affinities^12,13^ as well as designing *de novo* molecules satisfying a variety of constraints.^14,15^ The binding affinity of a given molecule to a protein target is a function of the positions and identities of its atoms, which can be approximated by a neural network in a machine learning-based scoring function (MLBSF). Given the coordinates of a protein-ligand complex (obtained by docking or otherwise), such a function considers all interactions between the two molecules. In this way, the interaction energy between two molecules can be predicted.

However, as described above, there are growing concerns regarding the generalisability of machine learning models. Indeed, recent research into MLBSFs has highlighted their tendency to learn dataset biases, rather than physical interactions.^5–7,16^ In these cases, the MLBSFs perform less of an assessment of atomic contributions, and something more analogous to a nearest-neighbours search in ligand-pocket space. This is useful if the application is to proteins or scaffolds with a large amount of previous binding data available, but not for new targets or ligands, as the true functional relationship governing binding is not learned. This problem is often exacerbated by a lack of rigorous dataset filtering. One of the most widely used benchmarks for scoring functions, both classical and machine learning-based, is the Comparative Assessment of Scoring Functions, 2016 (CASF-2016).^17^ The usual method of training a MLBSF is to train on the PDBBind^18^ general set of crystal structures and binding affinities after removing the structures in CASF-2016. As we found when building our scoring function, this approach suffers from information leakage concerning almost every test structure, resulting in an overestimation of the accuracy of MLBSFs.

There is an active area of research surrounding how machine learning models make predictions,^19–22^ a technique known as input attribution. This can be applied to MLBSFs, with the idea being that an ideal MLBSF—one which has learnt to distinguish atomic interactions, rather than just dataset biases—should be able to identify the most important binding interactions taking place in a given bound structure. This should mean that the function predicts well on unseen targets.

In practice, the type of attribution that can be used with an MLBSF depends on its architecture. The two prevalent deep learning architectures used as scoring functions are convolutional neural networks (CNNs)^23–25^ and graph neural networks (GNNs).^26–28^ Attribution for CNN-MLBSFs is limited, because spatial relationships are lost as information flows deeper into the network.^29^ Masking—a method by which a feature is allocated a score equal to the difference between the model’s prediction made when the feature is removed from the input, and the score when it is present— can still be used, but this is a computationally expensive method.

Another method for performing attribution is attention analysis. Attention mechanisms have occasionally been used in graph-based MLBSFs to encourage models to differentiate the contribution of each interaction to binding affinity.^28,30^ These work by allocating scores to the graph’s edges, which can be thought of as indicating how much importance is ascribed to each edge by the model. In contrast to CNNs, GNNs preserve the concept of the atom until the final layers of the network, so edges can be interrogated for information. This is helpful in instances where an attention mechanism has been used to weight the edges, as we gain direct insight into how important the network considers each edge to be. However, none of the current leading models for affinity prediction use attention layers for intermolecular interactions, so attribution cannot be carried out on them in this way.

If input attribution could be carried out on MLBSFs, it could be used to extract binding insights from a protein, for example, for fragment-based drug discovery (FBDD); a strategy that seeks to identify simple, small molecules (fragments) that interact with protein targets, and grow them into active leads. In order to meaningfully add atoms to these molecules, structural knowledge about the target needs to be considered. This can be extracted from known binders, but this limits the exploration of chemical space as well as restricting the applicability of the approach to targets with known ligands. Extracting structural information directly from the protein itself is therefore advantageous. Currently, there is only one available generative model for fragment elaboration that relies solely on protein structure.^31^ This method uses a data-driven approach^32^ to identify ‘hotspots’: regions in the protein pocket that contribute disproportionately to binding. Currently there exists no deep learningbased tool to identify such regions.

In order to have full control over the architecture and training set of the MLBSF we perform attribution on, we built PointVS, an *E*(*n*)-equivariant graph neural network. It is an MLBSF designed to predict both binding affinity and pose score. We assess its performance on the CASF-2016 benchmark, and find it achieves comparable results to other leading scoring functions in spite of the additional rigorous filtering of the training dataset. We then use attribution to show that the model successfully identifies binding interactions in agreement with a distance-based interaction profiler.^33^ To further investigate the power of the attribution method, we use it with fragment screen data to obtain information that can be leveraged to perform fragment elaboration. We see that using this information extraction method results in improved docking scores compared to using a data-driven approach. We make our code as well as our unbiased test and train splits available on GitHub at github.com/oxpig/PointVS.

## 2. Methods

An overall schema of the methods used to debias and test PointVS is shown in Fig 1.

**Fig. 1.**
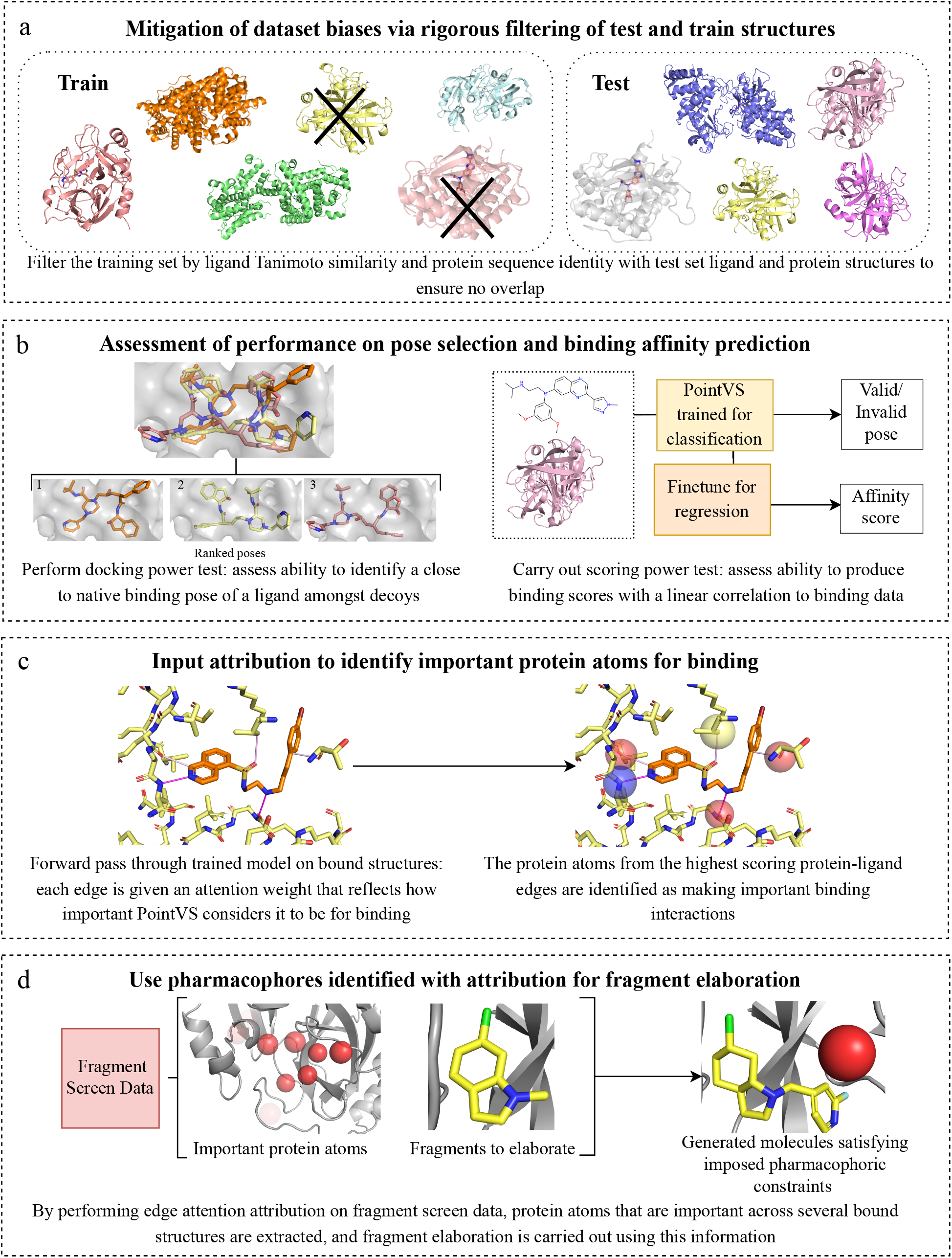
An overall schema of the methods used to debias and test PointVS. We first thoroughly filter the test and train sets (box a), before benchmarking the performance of PointVS on the docking power and scoring power tests (b). We then use attribution to gain insights into important binding regions in the protein pocket (c), which we use for fragment elaboration (d).

### 2.1 Building and Testing PointVS

#### 2.1.1 Model

PointVS is a lightweight *E*(*n*)-equivariant graph neural network layer model, consisting of an initial projection to take the number of node features from 12 to 32, then 48 EGNN layers, followed by a global average pooling of the final node embeddings. This is followed by a sigmoid layer which gives a label *y* ∈ [0, 1] during the first stage of training on pose prediction. During finetuning on affinity data, this final layer is replaced by a randomly initiated fully connected layer and ReLU activation, which outputs **y** ∈ (ℝ^+^)^3^ (see Fig 2). It also uses a shallow neural network as an attention mechanism,^34^ which learns to score network edges—in this case, representing atomic interactions—by their importance.

**Fig. 2.**
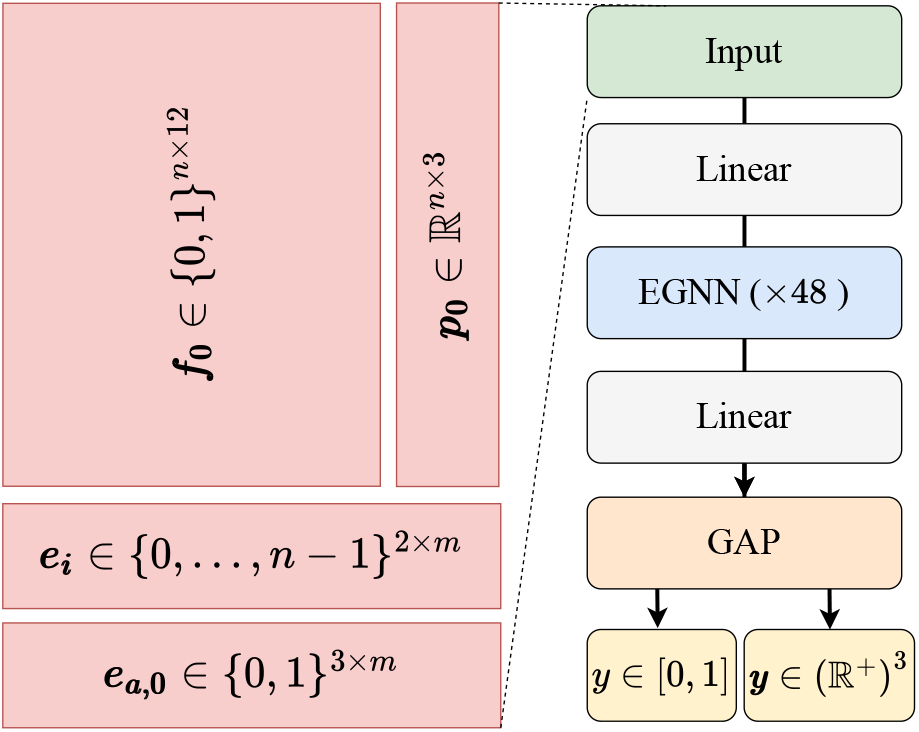
Architecture and input format of PointVS. Each of the *n* atoms in the input to the model (left) is given a one-hot encoded feature vector with a single bit added to indicate whether the atom is from the ligand or the receptor, as well as a position ***p***_**0**_ ∈ ℝ^3^. There are *m* edges, defined by the edge indices ***e***_***i***_ ∈{0, …,*n* −1 ^2 ×*m*^ which are the indices of connected atoms, and the corresponding edge attributes ***e***_***a***_ ∈{0, 1} ^3×*m*^. These are one-hot encodings representing ligand-ligand, ligand-protein, and protein protein edges. There are skip connections between each of the EGNN layers, and the linear and global average final pooling (GAP) layers only act upon the node features *f*.

#### 2.1.2 Datasets

A full list of the datasets used is given in Table 1, and more information about each of them is provided in the SI.

**Table 1.** Datasets, their sizes and the number of unique PDB structures in their generation. Redocked includes no structures with the same PDB code as any structures in the gninaSet_*Pose*_ or Core sets, and the General set shares no PDB codes with the Core set. In the Task/Split column, ‘C’ refers to pose classification and ‘R’ refers to affinity regression.

| Dataset | Size | Unique PDB IDs | Task/ Split |
| --- | --- | --- | --- |
| Redocked | 780,572 | 19,595 | C/Train |
| Redocked\gninaSet<sub>Pose80</sub> | 633,047 | 17,528 | C/Train |
| Redocked\Core<sub>80</sub> | 629,963 | 16,900 | C/Train |
| General | 19,157 | 19,157 | R/Train |
| General\Core<sub>80</sub> | 14,555 | 14,555 | R/Train |
| gninaSet<sub>Pose</sub> | 20,411 | 411 | C/Test |
| Core | 285 | 285 | R/Test |

Due to the lack of affinity data, we choose to pretrain on a pose classification task in order to learn some of the physics of binding before finetuning on affinity data.

Our training dataset for pose classification is Redocked2020^35^ (Redocked from hereonin), with the test set, gninaSet_*Pose*_, generated according to McNutt *et al*.^23^ so that classification performance can be directly compared with their work. This test is similar to the *docking power* test from CASF-16,^17^ which we also conduct.

For regression on binding affinity values, we train a pose classifier on the Redocked set, finetune on affinity data from the PDBBind General Set, and test on the PDBBind Core Set. This replicates the *scoring power* test from CASF-16. As the Core Set is a subset of the General Set, we remove Core Set structures from the training data.

Filtered subsets of both the Redocked set and the PDBBind v.2020 General Set are generated for both of these stages of training, as well as random subsets of the original training sets of the same size as the filtered sets. The filtered datasets were constructed by removing proteins and ligands if they met either of the following criteria:

1. Tanimoto similarity of the 2048-bit Morgan fingerprint between the ligand and any of the 285 test set ligands greater than 0.8

2. Sequence identity between the protein and any of the 285 test set proteins greater than 0.8

These filters were also applied to the Redocked set with respect to the gninaSet_*Pose*_ test set. All files which could not be parsed by either RDKit^36^ or OpenBabel^37^ were also discarded.

This process reduces the training set sizes from 780,572 to 633,047 and 629,963 (Redocked/gninaSet_*Pose*80_ and Redocked*/*Core_80_) and from 19,157 to 14,555 (General Core_80_). In order to deconvolute the effects of training set size from the effect of the filters, we also constructed training sets of the same size, randomly sampled from the original training sets. These are called Redocked\gninaSet_*PoseR*_, Redocked\Core_*R*_ and General Core_*R*_. Carrying out filtering in this way should ensure the PDBBind Core set performance is a more accurate reflection of accuracy on unseen targets and ligands.

#### 2.1.3 Performance Metrics

We carried out the pose ranking test as defined by the authors of gnina^23^ as well as the CASF-16 docking power and scoring power tests.

Docking power refers to a scoring function’s ability to identify a ligand’s native binding pose among decoys. It is quantified using Top-N, which is the percentage of systems with a “good” pose ranked in the top N. A pose is considered good if the RMSD to the crystal pose is less than 2Å.

The scoring power is defined by Su *et al*. as “the ability of a scoring function to produce binding scores in a linear correlation with experimental binding data”.^17^ This is best measured using the Pearson correlation coefficient (PCC) between the score given by the scoring function and the measured binding affinity data. For training our networks, we once again used both filtered and unfiltered versions of the PDBBind General dataset to test the influence of dataset bias.

### 2.2 Attribution

A number of previous studies have identified the tendency of MLBSFs to make predictions based on dataset biases rather than an understanding of the physics of binding: dataset clustering and cross-validation have been used to demonstrate that many CNN-based MLBSFs, for instance, are prone to distinguishing binders from non-binders based not on protein-ligand interactions, but on ligand features alone.^5–7,38^ Ensuring that predictive power is diminished in the absence of receptor information in the test set causes at least some receptor information to be used.^39^ However, only directly observing which parts of the input space are used in making predictions can give affirmative proof that machine learning models are learning to identify physical interactions that separate active from inactive poses.

Interpretability in machine learning for pose prediction and virtual screening is an ongoing problem; the architecture of most CNNs causes the concept of individual atoms to be lost as information flows deeper into the network, making them fundamentally unsuitable to attribution at the atomic level. By contrast, graph-based architectures such as PointVS maintain distinct nodes (atoms) until the final layers. These, having passed through the entire network, are rich in information and are directly related to the relevant atom in the input. As the node features are directly attributable to their input atom, and the final pose score is a function of the node features, there is a direct relationship between the atoms and the PointVS score, although there is no incentive during training to localise class contributions on particular atoms. The edges between atoms can also be probed for how much importance is ascribed to atomic interactions, as an intuitive way to describe non-covalent bonds. As set out in Fig 1, we can also use this knowledge of importance assigned by PointVS to edges to identify important protein atoms for binding.

To check whether PointVS has learnt to identify important binding interactions, and further, whether these can be extracted with attribution, we compare the results of performing attribution on PointVS to those found when using other leading MLBSFs.

#### 2.2.1 Attribution Techniques

We use three approaches for attribution: atom masking, bond masking, and edge attention.

##### Atom masking

Atom masking is a process by which the importance of different atoms can be ascertained. It is carried out by calculating the difference between the score given when an atom *i* is removed from the set of input atoms, *X*, and the score when it is present. As all architectures of MLBSFs take as input atom information, atom masking can be carried out on any MLBSF.

##### Bond masking

Similar to atom masking, bond masking is a process by which the importance of different bonds can be ascertained. It is carried out by calculating the difference between the score given when an edge *e* is removed from the input graph encoding the protein-ligand complex and the score when it is present. Whilst atom masking can be carried out with CNNs and GNNs alike, bond masking can only be carried out with architectures that take as input not only information about the atoms in the complex, but also information about the connections between atoms. Graph representations of molecules, which use nodes and node features to describe atoms, and edges and edge features to describe interactions between them, are well-suited to this type of attribution. CNNs, which take only atom information as input, are not.

##### Edge attention

GNN-based MLBSFs can be built to include an attention-like mechanism, where the edges are given a score between 0 and 1 by a shallow multilayer perceptron (MLP) which takes as input the edge embeddings themselves. These weights can be thought of as indicating how much each edge is weighted by the network, so attribution can easily be performed by extracting the edge attention weights.

In order to facilitate attribution, PointVS was built to include an attention-like mechanism. After the edges are assigned attention scores, the edge message passed to the *i*^*th*^ node at each layer **m**_*i*_ is the sum of connecting edge embed-dings **m**_*i j*_ in the neighbourhood of the node *N*(*i*), weighted by this attention score *e*_*i j*_, as in Equation 8 in Satorras *et*.*al*:^40^

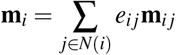

We carry out attribution with PointVS as well as two other MLBSFs for comparison: a convolution-based method, gnina,^23^ and another graph-based model, InteractionGraphNet.^28^

#### 2.2.2 Other MLBSFs gnina

Gnina^23,41^ is one of the leading tools for predicting protein-ligand binding, and a popular convolution-based tool. It consists of an ensemble of CNNs, which take as input a bound structure, and output either a pose score or binding affinity.

In recognition of the difficulty associated with performing attribution with convolution-based scoring functions, the authors published a paper describing various methods that can be used to carry out attribution with gnina.^29^ The code implementing these methods, gninavis, is currently unavailable, so we instead implemented a standard atomic masking procedure. This was carried out by generating a PDB file for each atom in the bound structure, from which said atom was removed. These were all scored by gnina, and their respective results compared to the score of the complete PDB.

##### InteractionGraphNet

We also carried out bond masking on InteractionGraphNet (IGN).^28^ This uses two graphs—one intramolecular and one intermolecular—to describe the bound structure. The intramolecular graph uses an attention mechanism similar to that of PointVS to ascribe weights to edges, but as this is only applied to edges within the same molecule, these weights cannot be probed for information about binding interactions.

The authors of the paper highlighted that IGN’s architecture was chosen to try and force the model to learn the key features of protein–ligand interactions rather than dataset bias. However, a recent paper that performed clustering and cross-validation on an array of MLBSFs^42^ showed that although IGN was one of the best predictors when assessed on a conventional test set, it saw a substantial decrease in performance when cross-validation was carried out. The scoring power PCC (defined above) decreased from 0.80 on the default test set to 0.47 when a pocket similarity-based clustering (a process that ensured that no bound structures with similar pocket structures were in both the train and test sets) was used. This decrease in performance is a sign of IGN making predictions based on dataset biases rather than an understanding of the physics of binding.

We performed bond masking attribution on IGN as follows. First, we supplied a bound structure to the model. The two graph types—intramolecular and intermolecular—arere then constructed. Edges in the intermolecular are initialised if they are found to connect atoms that are less than 8Å apart. We then perform masking on any edge that joins atoms at a distance of less than 4Å (for consistency with PointVS) by generating a new graph with said edge deleted. These are then passed through the network, and their scores compared to the score of the full graph.

### 2.3 Fragment Elaboration

To elaborate a fragment towards more potent, lead-like compounds, information about important areas of the protein pocket being targeted is key. We set up a series of tests to investigate if the attribution scores from PointVS can identify these important sites.

By performing attribution on a crystal structure of a given protein with a small molecule bound, we obtain binding information in the form of an importance score for every protein atom less than 6Å away from any ligand atom. If a fragment screen has been carried out on said target, the attribution process can then be repeated for several crystal structures to obtain an average importance score for each of the protein atoms. We use this list of protein atoms and associated scores as hotspots to perform fragment elaboration. To assess them, we say that the quality of a hotspot—how important the point it is placed on actually is for binding— is proportional to the binding affinity of the molecules generated in the elaboration process, which we assess using a metric derived from docking score, ΔSLE_*α*_.

We compare the ability of PointVS hotspots to highlight important binding regions to traditional, data-based fragment hotspot maps. To obtain these, we use the Hotspots API,^32^ which implements the algorithm described in Radoux *et al*.^43^ We then process the output of the API as in Hadfield *et al*.^31^ A description of this process is given in the SI.

For each target we perform fragment elaboration tests on, we obtain hotspot maps with PointVS using the crystal structures of their fragment screens available on the fragalysis platform.^44^ The structures used are listed in SI Table 1. To obtain traditional hotspot maps, we run the Hotspots API on one structure of the target randomly selected from the fragalysis set of bound structures.

We also use the fragalysis set of crystal ligand structures listed in SI Table 1 to obtain fragments to elaborate. To do this, we enumerate all cuts of acyclic single bonds that are not part of functional groups, and add a ‘dummy’ atom at the site of the cut. This atom is where atoms will be added to the fragment.

To perform fragment elaboration using the two sets of hotspot maps, we use STRIFE, a generative model for fragment elaboration. We follow the method outlined for its use in Hadfield *et al*.^31^ We take as input a fragment, a specified exit atom, and information about one hotspot; namely, its coordinates and type. STRIFE then generates molecules matching the pharmacophoric profile specified.

To achieve this, STRIFE has two main stages. The first of these, exploration, aims to generate a set of elaborations that contain an acceptor or donor in close proximity to a hotspot. These elaborations are then docked using GOLD’s constrained docking functionality,^10^ and those which successfully place a functional group within a certain distance (2Å for the API hotspots, which are in the protein pocket, and 3Å for PointVS hotspots, which are on protein atoms) of the hotspot being targeted are selected as ‘quasi-actives’. In the next stage, refinement, these quasi-actives are used to derive a fine-grained pharmacophoric profile. STRIFE then generates elaborations using those pharmacophoric profiles, and docks them. These docked elaborations constitute its final output.

For a given hotspot, we perform elaboration on a set of fragments. The size of this test set depends on the number of bound structures of ligands available for the target in question (see SI Table 1). For some fragment-hotspot pairings, elaborations cannot be generated. This can be due to the hotspot in question being too close or too far from the exit point on the fragment, or at an angle that is unfavourable to elaborate along. In these cases, the generation stage will be unable to produce any quasi-actives; in other words, it will be unable to generate any molecules that, when docked, have a hydrogen bond acceptor or donor within a certain distance from the hotspot. An example of this is shown in Table 5, which shows the number of fragments successfully elaborated in the pocket of Mpro for every hotspot tested. We see that the hotspot ranked ninth by the Hotspots API, for instance, cannot be reached by elaborating on any of the 109 fragments tested, and hence no molecules are output.

The number of final elaborations generated is dependent on the number of quasi-actives that the first stage, exploration, generates. Although STRIFE is asked to generate 250 elaborations per fragment, there are often fragment-hotspot pairings where the number of quasi-actives identified is insufficient for the refinement phase to produce 250 distinct molecules. The mean number of elaborations generated for a given pair is then less than 250, as seen in Table 5. Nevertheless, as each hotspot is tested on a number of fragments occupying various positions in the pocket, we still obtain a large number of generated molecules for a given hotspot.

To assess the generated molecules, we dock them using GOLD,^10^ the CCDC’s protein-ligand docking software. We use the constrained docking functionality—constraining using the fragment, for which we have a crystal structure— from which each molecule receives a score. We then calculate a ligand efficiency by dividing this score by the number of heavy atoms present in the molecule. This is to compensate for docking algorithms tendencies to favour larger molecules. From the ligand efficiency, we derive a standardised ligand efficiency. This is obtained by standardising the ligand efficiencies of the generated molecules and the ground truth molecule for given fragment-hotspot combination to have zero mean and unit variance (here, ground truth molecule refers to the molecule that was segmented to obtain the test fragment). We then sort the standardized ligand efficiencies of the elaborations from highest to lowest scoring, and take the mean of the top *α*. In this work, we use *α*=20. Subtracting the ground truth ligand efficiency value from this value then provides us with ΔSLE_20_ for every fragment-hotspot pair, so for a given hotspot, we take the mean over the ΔSLE_20_s of all the fragments successfully elaborated towards it. We use this metric as it provides an insight into how the elaboration process when using a particular hotspot has impacted the original molecule’s docking score: a positive ΔSLE_*α*_ means the top elaborations are improvements on the original molecule, and a negative ΔSLE_*α*_ shows that the elaboration process has decreased the ligand efficiency with respect to the starting molecule.

## 3. Results

We tested the ability of our MLBSF, PointVS, to perform pose selection and affinity prediction. Through the use of attribution, we also verified not only that PointVS has successfully learnt to recognise binding interactions, but that when given a number of bound structures of a given target, it could be used to extract information about important binding pharmacophores that could then be used in automated fragment elaboration.

### 3.1 Training and Testing PointVS

#### 3.1.1 Bias in CASF-16

We implemented a rigorous filtering step to ensure that the training set and test set structures used by PointVS have no overlap (see methods). In order to compare PointVS with other machine learning methods which have not included this filtering step,^23,45,46^ we also constructed a smaller test set out of the PDBBind Core set by applying the same filters, but excluding test set structures rather than train set structures. However, we found that such a test set would only contain a single structure: 284 of 285 of the Core set proteins have a counterpart with 90% sequence similarity in the General set. Further, 273 of them (95.7%) have a counterpart with an identical sequence, making any fair and unbiased comparison to these methods impossible.

#### 3.1.2 Docking and scoring power tests

We carried out the pose ranking test as defined by the authors of gnina^23^ as well as the CASF-16 docking power and scoring power tests. The results of these tests are shown in Fig 3, Table 2, and Table 3.

**Table 2.** Top-1 pose ranking performance for different models and training sets (in brackets) on the PDBBind Core set (the CASF-16 docking power test set). The non-machine learning scoring function is shown in italic. The performance is shown for both the case where the crystal structure of the ligand is included in the set of poses being ranked, and the case where it is not included, in which case the native pose is defined as any pose less than 2Å RMSD away from the crystal pose. The highest Top-1 for each case is shown in bold. Not included in the table are ΔVinaXGB and ΔVinaRF, for which only the performances on the crystal pose included test are provided by the authors as 92 and 90 respectively.

| Model | Top-1 |  |
| --- | --- | --- |
|  | Crystal pose included | Crystal pose not included |
| PointVS (Redocked\Core<sub>R</sub>) | 90.5 | 84.9 |
| PointVS (Redocked\Core<sub>80</sub>) | 91.2 | 84.2 |
| PointVS (Redocked) | **91.3** | **85.3** |
| gnina (Redocked) | 91.2 | 83.2 |
| *Autodock Vina* | 90.2 | 84.6 |

**Table 3.** Pearson correlation coefficients between measured and predicted affinity for different scoring functions on the PDBBind Core set. Scoring functions are split into biased and debiased methods according to the overlap between their training sets, which are shown in brackets where applicable, and the Core set. Non-machine learning scoring functions are shown in italic. The highest PCCs obtained with both biased and debiased methods are shown in bold.

|  | Scoring Function | PCC |
| --- | --- | --- |
| Biased | gnina (Redocked) | 0.753 ± 0.008 |
|  | gnina (General) | **0.816 ± 0.008** |
|  | PointVS (General) | 0.805 ± 0.010 |
|  | PointVS (General\Core<sub>R</sub>) | 0.803 ± 0.012 |
|  | ΔVinaXGB | 0.796 |
|  | ΔVinaRF | 0.732 |
| Debiased | PointVS (General\Core<sub>80</sub>) | **0.754 ± 0.015** |
|  | *X-Score* | 0.631 |
|  | *Autodock Vina* | 0.601 |

**Fig. 3.**
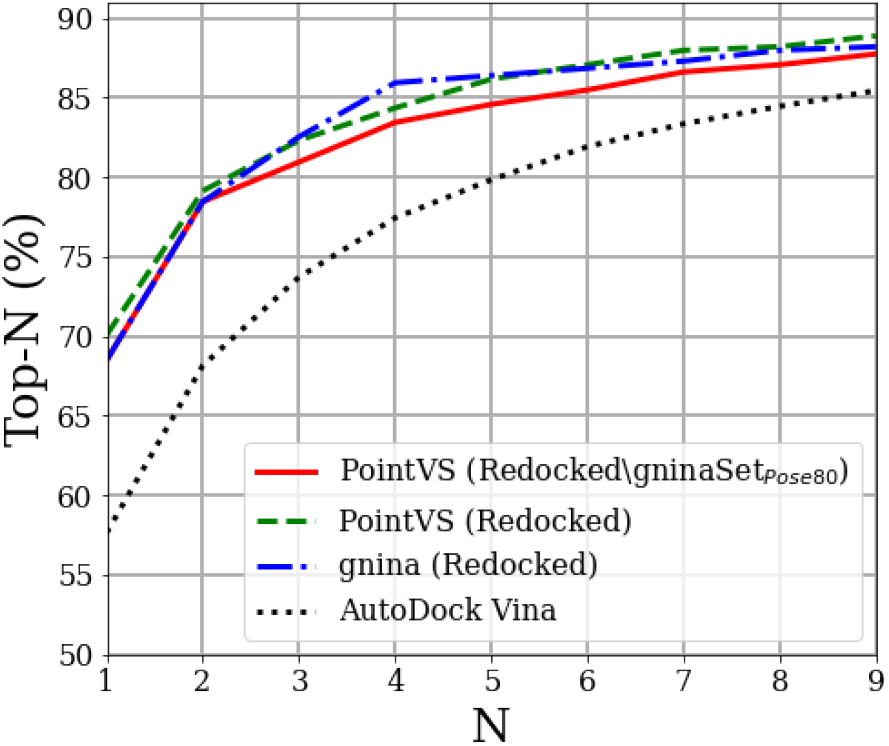
Top-N vs N pose ranking performance on the gninaSet_*Pose*_ set for Autodock Vina, PointVS and gnina. Terms in brackets in the legend refer to the training sets; filtering the training set by protein and ligand similarity results in slightly degraded performance. The Top-1 values for PointVS trained on Redocked\gninaSet_*Pose*80_ and gnina trained on Redocked are the same (68%), with PointVS trained on Redocked achieving 70%.

Top-N pose ranking performance on the gninaSet_*Pose*_ set is shown in Fig 3. When the training sets are the same, PointVS ranks a good pose at the top in 70% of cases, as opposed to 68% with gnina. When PointVS is trained on the smaller Redocked\gninaSet_*Pose*80_ set, such that no similar proteins or ligands as the test set are seen during training, this drops to 68%. This suggests that for pose prediction, filtering the dataset to remove overlap between train and test structures only causes a minor decrease in docking power.

The results of the CASF-16 docking power test (Table 2) support this conclusion: the difference in performance between all of the methods is minimal, and is not affected significantly by whether or not the training data has been filtered for structural similarity.

On the scoring power test, the three best methods by scoring without dataset filtering (gnina, PointVS and ΔVinaXGB) achieve very similar performance, and PointVS outperforms the best non-machine-learning methods from the original CASF-16 work,^17^ Autodock Vina, significantly (0.805 vs 0.601). However, the performance of PointVS when trained on the filtered dataset General\Core_80_ is a better indicator of its true performance on unseen protein-ligand combinations with no analogues in the training data.

To test whether the drop in the performance of PointVS when similar structures are removed from the training set is related to bias or change in training data size, we trained PointVS on General\Core_*R*_. This has the same training set size as General\Core_80_, but can still contain similar structures to those in the test set. PointVS trained on the General\Core_*R*_ achieves a PCC of 0.803 on the CASF-16 set. This is almost identical to the PCC of PointVS trained on the entire General set (PCC of 0.805), and significantly higher than PointVS trained on the General\Core_80_ set (PCC of 0.754). This points to the performance of MLBSFs when trained and tested using biased datasets being boosted, and contrasts with our findings in pose prediction performance, in which we saw minimal change when filtering was performed.

The three machine learning methods which use three distinct featurisation methods and three separate architectures (PointVS, gnina and ΔVinaXGB) obtain similar PCCs when trained on the unfiltered General set, suggesting that the limiting factor in predictive performance is what information the training data holds about the test set data, rather than the architecture or featurisation. PointVS does not outperform other leading scoring functions: in fact, there exist several scoring functions which report higher performance for the tests carried out.^47–49^ However, given our findings surrounding the data leakage present in the CASF-16 tests, we suggest that performances reported for scoring functions trained on the unfiltered dataset are overly optimistic. Without the other leading models being retrained on the filtered dataset and their performances reevaluated, there is no way to provide a fair and unbiased comparison to them.

Furthermore, although our dataset filtering has gone some way to reducing the bias that PointVS learns, it is unlikely that we have successfully removed all dependence on it. Recent research has shown that sequence-based filtering for assessing model generalisability is flawed, as too much attention is paid to the entire length of the protein rather than the pocket itself.^42^ It is therefore likely that even our rigorously filtered datasets have some leakage, and that PointVS has still learnt some unintentional biases. Even the performance of the debiased model (particularly on the scoring power test, which appears to be more influenced by bias) is probably overly optimistic.

PointVS appears to perform at a similar level to leading scoring functions even when trained and tested using a less biased setup. In the next sections of the paper we take advantage of the fact that we can examine its edge attention scores in order to investigate if it has learnt about binding interactions.

### 3.2 Attribution: Identifying Important Binding Sites

PointVS appears to offer a substantial improvement in predictive performance of binding affinity over non-machinelearning methods. However, MLBSFs are notoriously difficult to interpret. Good performance when trained on debiased training data is circumstantial evidence that the interatomic interactions involved in binding are being learned, but demonstrating that the attribution scores of PointVS relate to important binding interactions is a more direct measure. We next present several results demonstrating that important interatomic interactions have been learnt by PointVS.

#### 3.2.1 Human Tankyrase-2 Inhibitors

The Protein-Ligand Interaction Profiler (PLIP)^33^ uses simple geometric rules to predict interactions at the interface between a protein and a ligand. We used PLIP to identify the “most important” bonds for three human Tankyrase-2 inhibitors, and examined whether the atoms forming these bonds were highlighted when attribution was carried out on gnina,^23^ IGN,^28^ and PointVS.

The structures of three tankyrase inhibitors are shown in Fig 4. The top row (structures a, b and c) shows the PLIP analysis of each structure. The structures in the other three rows show the results of attribution with gnina (second row, structures d, e and f), IGN (third row, structures g, h and i), and PointVS (bottom row, structures j, k and l). The structures in the second row are coloured by the scores obtained from atom masking with gnina, with atoms contributing negatively to binding being shown in red (negative score), neutral atoms in white, and positively-contributing atoms in green (positive score). Attribution with IGN and PointVS— which was carried out with bond masking and edge attention analysis, respectively—resulted in scores corresponding to interactions between pairs of atoms rather than individual atoms. The five highest-scoring edges for structures using both of these methods are shown in the bottom two rows, with dark red lines representing high scores from IGN in structures g, h and i, and darker pink representing higher scores from PointVS in structures j, k and l.

**Fig. 4.**
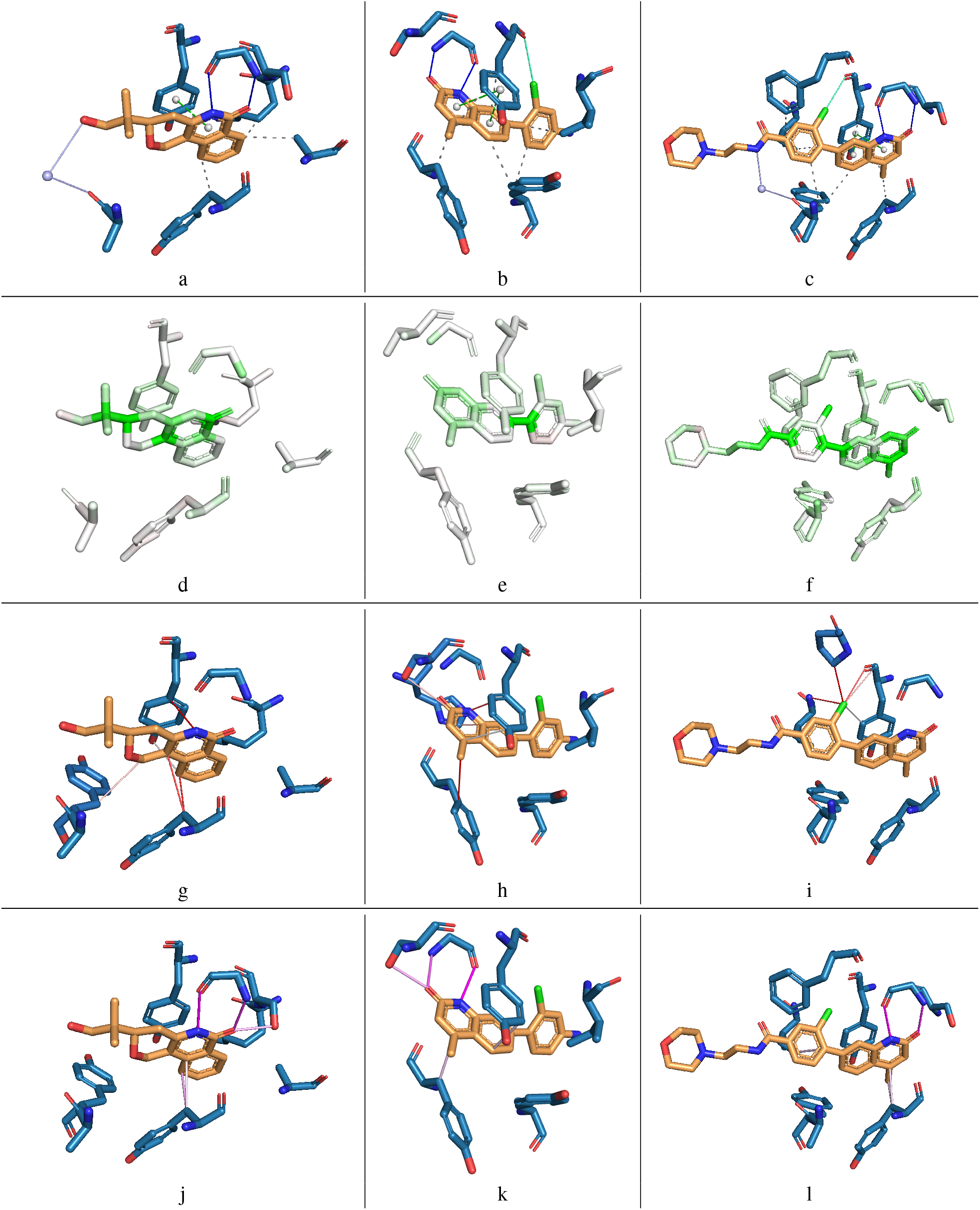
Three Tankyrase-2 inhibitors: 5C5P (a, d, g, j), 4J21 (b, e, h, k), and 4J22 (c, f, i, l). The top row shows the Protein-Ligand Interaction Profiler (PLIP) analysis of each structure, where dark blue lines are hydrogen bonds, dotted grey are hydrophobic interactions, dotted green are *π*-*π* interactions, solid green are halogen bonds, and lilac are water bridges. The second row shows the results of performing atom masking with gnina, with green representing a positive attribution score (identified as making a positive contribution to binding) and red a negative score (making a negative contribution). The third row shows the results of bond masking with IGN, with lines between atoms showing the top five highest-scoring edges for 11 each structure, with darker red representing higher scores. The bottom row shows the results of edge-attention attribution performed with PointVS, with the lines also showing the top five highest-scoring edges, and darker pink representing higher scores.

Considering the first column, the two ligand atoms that form hydrogen bonds with the protein as calculated by PLIP (Fig 4a) are scored by gnina as the 6th and 8th most important of 20 (Fig 4d). Of the two hydrogen bonding protein atoms, gnina recognises one of them as being extremely important (the highest scoring protein atom), and the other as neutral. The second structure, Fig 4e, shows a similar story, with the three important ligand atoms as given by PLIP (Fig 4b) scoring 4th, 7th and 8th out of 20, but the protein atoms they respectively bind to scoring negatively, neutrally and very slightly positively. In the third structure, the three bound ligand atoms as identified by PLIP (Fig 4c) are scored 11th, 13th and 14th of 30 ligand atoms, with the corresponding protein atoms scoring negatively, weakly positively and moderately positively (Fig 4f). Interestingly, the last two structures both contain halogen bonds, and in both cases gnina ranks the ligand atom as important, and the protein atom as a negative contribution. Overall, gnina struggles to identify important individual atoms (both in the ligand and protein) for binding. The atoms being scored individually rather than by pairs as with bond masking or edge attention (see below) makes it more difficult to identify the bonds that are being formed: of the bonds identified by PLIP, only in the first structure was there an example where both the protein and ligand atoms were given a high score by gnina, making it the only case where an important bond was identified.

When we performed atom masking on PointVS, we found that the results were similar to those from gnina, in that the atoms contributing to bonds were scored highly but not necessarily enough to recognise their true importance. However, as described below, we found that edge-based attribution methods provided a clearer insight into important interactions.

The first of the results achieved with edge-based attribution methods is the third row of Fig 4, which shows the edges found to be important when carrying out bond masking with IGN. In the first structure, Fig 4g, the bond highlighted as most important connects a ligand atom that PLIP highlights as forming a hydrogen bond (Fig 4a) with a different protein atom. At first glance, it appears that IGN has incorrectly predicted the formation of a hydrogen bond between these atoms, but it should be noted that the atoms that this edge connects are both in aromatic rings that PLIP identifies as being involved in a *π*-stacking interaction. IGN also correctly identifies a hydrophobic interaction, which is ranked as the second most important edge. In the second structure (Fig 4h), we again see edges connecting rings involved in *π*-stacking in Fig 4b, this time ranked first and fifth most important. However, none of the other interactions shown to be taking place in 4b—neither the hydrogen nor the halogen bonds—are recognised. This contrasts the third structure, 4i, which shows that IGN has identified the ligand atom that forms a halogen bond as extremely important, with all five of the highest-scoring edges connecting it to various protein atoms. The edge that connects it to the oxygen atom that PLIP identifies as being the other contributor to this bond (Fig 4c) is ranked third most important. In contrast to the first two structures, little importance is ascribed to the edges connecting the aromatic rings involved in *π*-stacking.

Altogether, the edges identified as important by performing bond masking with IGN show some agreement with PLIP. This is especially true of *π*-stacking interactions, of which it identifies two of three, and the least true of hydrogen bonds, of which it identifies none. In all three structures, IGN appears to successfully identify one of the interactions taking place (*π*-stacking in the first two, g and h, and halogen bonding in the third, i) but fails to identify any beyond those.

Considering the first example in the final row of Fig 4, we see that the bonds ranked first and second most important by PointVS (Fig 4j) correspond to the hydrogen bonds identified by PLIP (Fig 4a). Edges between the two aromatic rings involved in *π*-stacking are also given high importance scores, being ranked fourth and fifth most important. We see a similar pattern for the second structure, where the atoms involved in hydrogen bonds as defined by PLIP (Fig 4b) are connected by edges assigned high importance by PointVS (ranked first and second most important, Fig 4k). We also again see high scores assigned to the edges connecting the rings involved in *π*-stacking. PointVS does, however, fail to recognise the edge joining the atoms involved in the halogen bond shown in green in Fig 4b. In the third example structure, PointVS also identifies the hydrogen bonds identified by PLIP (Fig 4c) as being important, again ranking them as the most and second most important edges (Fig 4l). The other edges assigned high importance correspond to hydrophobic interactions.

The results above suggest that of the attribution methods tested, the two edge-based approaches are more effective at identifying important interactions taking place. However, of the two tested, only edge attention analysis with PointVS was able to reliably identify more than one of the bonds that PLIP identified as forming. This highlights that edge attention is a useful method for attribution, but further, that PointVS has gone some way to learning to identify important binding interactions.

#### 3.2.2 Large Scale Attribution Tests

To further verify that performing attribution with PointVS results in similar bonds being highlighted to those specified by PLIP, we performed attribution on the PDBBind Core set (the CASF-16 test set). For each bound structure, we found the top ten highest-scoring protein atoms (corresponding to the protein atoms connected to the ten highest-scoring edges). We then calculated the Spearman’s rank correlation, *ρ*, between the scores assigned to the top five and top ten scoring protein atoms, and the distance between the protein atom and the corresponding ligand atom. This results in a *ρ* value of 0.719 for the top five protein atoms, and a *ρ* value of 0.788 for the top ten protein atoms. This correlation implies that protein-ligand atom pairs that are close together (which in turn PLIP will consider more likely to be forming a bond) are commonly identified by PointVS. To ensure that this test is an effective way to learn how much a model bases its predictions on an understanding of binding rather than learnt biases, we carried out the same process using the biased PointVS model, and obtained a *ρ* value of 0.534 for the top five protein atoms, and a *ρ* value of 0.583 for the top ten protein atoms. This decrease in performance suggests that this test effectively measures the impact bias has had on the predictions made by a model.

We carried out the same process for IGN with bond masking and gnina with atom masking, but as both of these are significantly more computationally challenging than edge attention analysis, we used only 20 randomly selected structures from the Core set (see SI for list of PDB IDs). We also performed attribution with PointVS on this subset for comparison. The results of calculating both the rank correlation for the top five atoms, *ρ*_5_, and the top ten protein atoms, *ρ*_10_, are shown in Table 4.

**Table 4.** Mean rank correlation calculated between the scores of the top five (*ρ*_5_) and top ten (*ρ*_10_) ranked protein atoms by PointVS, gnina, and InteractionGraphNet (IGN) and the distance between them and the nearest polar ligand atom. The means were calculated over a subset of 20 randomly selected bound structures from the PDBBind Core set (see SI for PDB IDs).

| | $\rho_5$ | $\rho_{10}$ |
| --- | --- | --- |
| PointVS | 0.640 | 0.788 |
| gnina | 0.234 | 0.093 |
| IGN | 0.209 | -0.071 |

On this smaller set, we again see a correlation between PointVS scores and distance. In contrast, we see a much weaker correlation when using gnina or IGN (table 4). To assess the extent to which this can be attributed to the different attribution methods, we also carried out atom and bond masking with PointVS (see SI), and saw a decrease in performance compared to using edge attention.

These results once again suggest that PointVS is capable of identifying bonds that at minimum make sense geometrically, providing further evidence that the edge attribution scores from PointVS are identifying important binding interactions.

### 3.3 Hotspot Identification using PointVS

Having shown that PointVS is potentially capable of identifying important binding interactions, we next assessed whether it can be used as a method for extracting structural information from a number of bound structures, and hence used to guide molecule design. To do this, we extracted hotspot maps from PointVS by performing edge attention attribution on fragment screen data, and compared the resulting hotspots to those found using a data-driven approach, the Hotspots API.^32^

Our first test used the non-covalent bound structures available for the SARS-CoV-2 main protease (Mpro). The dataset available for this target is far more extensive than most targets will benefit from, as it is comprised of not only fragment-like molecules, but also follow-up compounds. These follow-up compounds, which have been curated with the specific intention of targetting important protein atoms, consist of closely related compounds with the same basic scaffold. Intuitively, the presence of highquality compounds should result in PointVS being able to extract higher-quality hotspots, as the bound ligands were designed to interact with protein atoms that have been experimentally validated to be important. We used this dataset as a starting point to verify that, given a selection of bound structures, PointVS can be used to identify these important protein atoms in a way that is useful for fragment elaboration.

The data was extracted from the fragalysis platform.^50^ A full list detailing the structures used is given in SI Table 1. The hotspots identified by the Hotspots API and attribution with PointVS are shown in Fig 5a and b. The hotspots of the two methods will not fall on exactly the same points, as the PointVS hotspots correspond to protein atoms, and the API hotspots to points in the binding pocket. Still, the two methods clearly highlight very separate regions of the binding pocket as important.

**Fig. 5.**
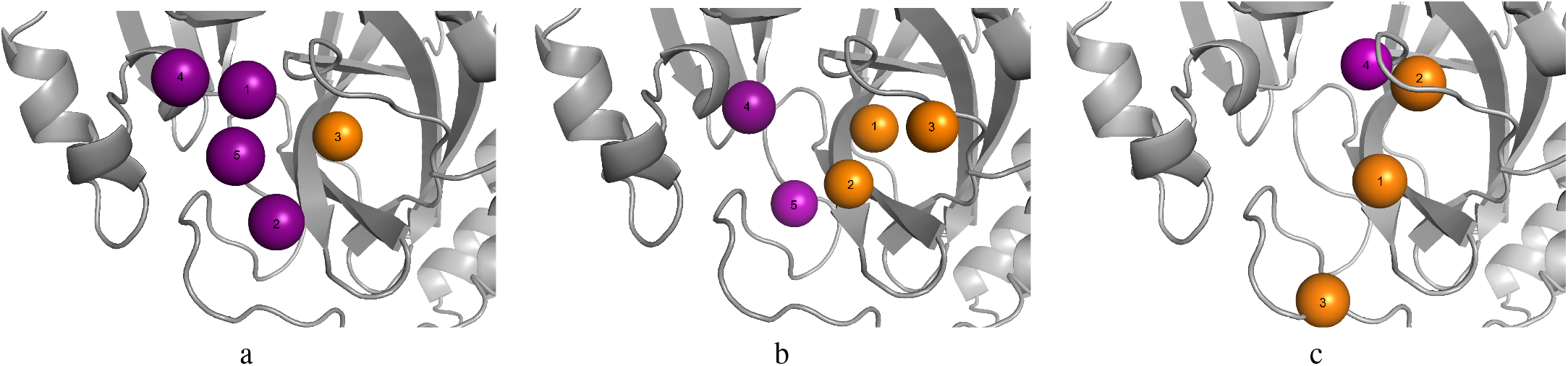
Donor and acceptor hotspot maps in the binding pocket of Mpro, coloured purple and orange respectively, and numbered according to their rank. The hotspots on the left (a) were obtained with the Hotspots API, the hotspots in the centre (b) were obtained with PointVS by performing attribution on 152 structures of bound fragments, and the hotspots on the right (c) were obtained by extracting the most common protein atoms identified by PLIP as involved in hydrogen bonds with ligand atoms across the 152 bound structures (in this case identified over five times).

#### 3.3.1 Impact of Fragment Screen Similarity

As mentioned previously, the Mpro dataset contains a significant number of closely-related follow-up compounds with the same basic scaffold. To ensure that PointVS is providing a novel insight into important binding regions in the pocket of Mpro and not highlighting important protein atoms by simply counting the number of times they are within close proximity of a ligand donor or acceptor atom, we used PLIP—which does use simple geometric rules to predict interactions—to extract the protein atoms that could be forming hydrogen bonds with ligand atoms for every bound structure. We then defined any protein atom that was highlighted more than five times across the 152 bound structures as a PLIP hotspot. This cutoff of five interactions resulted in only four hotspots being extracted. These top four are shown in Fig 5c, and correspond to protein atoms that were highlighted by PLIP 146, 40, 9 and 5 times out of the possible 152.

All but one of the PLIP hotspots are within 2Å of one of the top five PointVS hotspots. However, the top-ranking PLIP hotspot (which corresponds to a protein atom that forms a hydrogen bond in 146 of the 152 bound structures) is ranked second by PointVS, and the second PLIP hotspot (which PLIP identifies as interacting with ligand atoms in 40 structures) is ranked top by PointVS. As the first PLIP hotspot is at most the length of one hydrogen bond away from a ligand acceptor in almost every structure, we would expect a geometry-based method to clearly identify it as the top hotspot, as PLIP does. This suggests that PointVS is identifying important protein atoms with a more nuanced approach than simply counting interactions identified with geometric rules. These results do not mean the PointVS hotspot extraction method is immune to being biased when supplied with fragment screens containing clusters of ligand donor and acceptor atoms—however, it is a promising sign that it is not entirely biased by such clusters, and that PointVS is able to identify other areas in the pocket as important. This is a useful finding in light of recent research into fragment screen libraries has highlighted that even screens designed to be diverse structurally offer a limited exploration of protein pockets.^51^

#### 3.3.2 Dependency on Fragment Screen Size

Given that PointVS identifies sites that are different from a traditional hotspot method and a geometry-based method, we next tested whether the high-scoring sites identified by PointVS are dependent on the size of the input fragment set.

To do this, we performed the same hotspot extraction process as before, but instead of using all the fragment screen data (SI Table 1) at once, we randomly sampled smaller sets from it. We varied the size of the set sampled between 10 and 80. We then took the top five highest-scoring hotspots generated using PointVS with each randomly sampled set of fragment hits. The results of this are shown in Fig 6, where we took samples of size 10, 30 and 80 from the available bound structures (further images using other dataset sizes are given in SI Fig 2). This random sampling process was carried out 40 times. We see that when the hotspot maps are extracted from a smaller number of bound structures, a wider range of protein atoms are identified as being important; PointVS is less able to consistently pick out the same five protein atoms every time, and increasing the number of bound structures given to PointVS results in more consistent hotspot maps being generated. Nevertheless, even when using the smallest datasets containing only 10 bound structures, the five hotspots that are identified when using the full fragment screen are the most commonly included, hence their darker colour in Fig 6. This suggests that attribution with PointVS could also be a useful tool for less

**Fig. 6.**
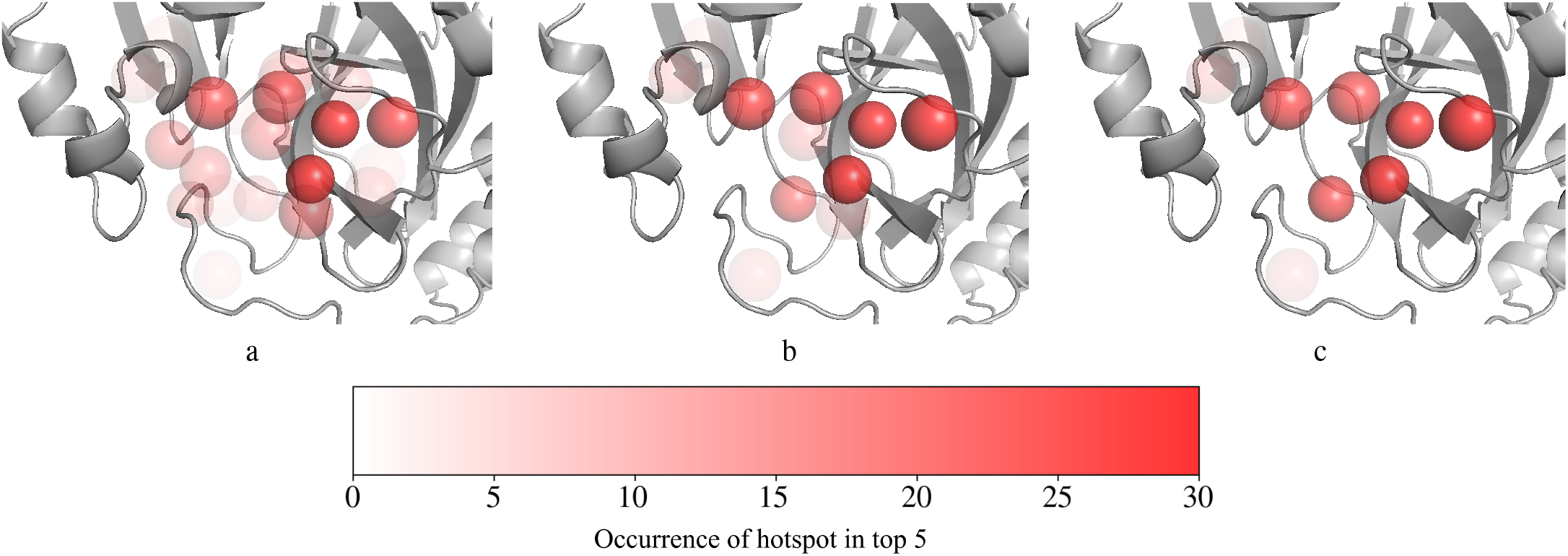
Top five scoring hotspot maps in the binding pocket of Mpro found by performing attribution on 40 sets of randomly sampled sets of 10 (a), 30 (b) and 80 (c) bound structures. Both donor and acceptor hotspots are shown in red. The spheres representing the hotspots are transparent and overlaid, so the more opaque a hotspot appears, the more times it was ranked in the top five.

### 3.4 Fragment Elaboration

Having demonstrated that the PointVS hotspots are consistent amongst themselves, and that they highlight different areas in the Mpro binding site to the API hotspots, we assessed whether they correctly highlight important areas by performing fragment elaboration. If PointVS is successfully identifying important interactions, molecules generated to satisfy the pharmacophoric constraints imposed by PointVS hotspots should be as good, if not better, than molecules generated to satisfy the API hotspots.

We performed attribution on the full set of Mpro bound crystal structures (see SI) to obtain PointVS hotspot maps, and used the Hotspots API to obtain a second set of hotspot maps. We used each of the top ten ranked hotspots from each set of maps to elaborate on 109 fragments using each of the top ten ranked hotspots from each set of maps (see methods). The number of fragments that were successfully elaborated—referring to cases where STRIFE^31^ was successfully able to grow towards the hotspot, and the donor or acceptor atom placed remained near to the hotspot after docking—using each hotspot is shown in Table 5, alongside the mean number of elaborations made per fragment.

**Table 5.** The number of fragments that were successfully elaborated on using PointVS and API hotspots for Mpro, and the mean number of elaborations that were generated for each fragment. STRIFE was provided with 109 fragments to elaborate on for each hotspot, and asked to generate 250 elaborations for each one.

| Rank | PointVS |  | Hotspots API |  |
| --- | --- | --- | --- | --- |
|  | Fragments successfully elaborated | Elaborations per fragment | Fragments successfully elaborated | Elaborations per fragment |
| 1 | 57 | 215 | 26 | 189 |
| 2 | 82 | 182 | 57 | 183 |
| 3 | 51 | 202 | 59 | 207 |
| 4 | 14 | 199 | 9 | 158 |
| 5 | 21 | 223 | 28 | 179 |
| 6 | 60 | 183 | 23 | 162 |
| 7 | 46 | 228 | 3 | 220 |
| 8 | 78 | 216 | 54 | 197 |
| 9 | 24 | 215 | 0 | 0 |
| 10 | 47 | 218 | 28 | 179 |

To assess the binding affinity of the generated elaborations, we docked them, as well as the fragment they were grown from, and used both docking scores to calculate a standardised ligand efficiency score, ΔSLE. This is the difference in ligand efficiency between the starting fragment and the elaborated compound (see methods). We then ranked the elaborations by these scores, and took the average of the top 20 scores to obtain ΔSLE_20_. To see the performance of a hotspot over all fragments successfully elaborated with it, we took the mean of the ΔSLE_20_s obtained.

The highest scoring elaborations by our ΔSLE_20_ metric were generated using the top five PointVS hotspots (Table 6). Elaborations made using the hotspots ranked 6-10 by PointVS have worse ΔSLE_20_s. By contrast, the Hotspots API successfully identified ten high-scoring hotspots (indeed, on average, these top ten hotspots produced elaborations with higher standardised ligand efficiencies than the PointVS hotspots), but was unable to successfully rank them amongst themselves. The accurate ranking of hotspots is important, as in real-world fragment-to-lead campaigns, only a small number of them can easily be targeted.

**Table 6.** The mean standardised ligand efficiency of the top 20 highest-scoring molecules (ΔSLE_20_) generated using hotspots ranked 1-5 and 6-10 by PointVS and the Hotspots API for Mpro. The highest ΔSLE_20_ for both ranges of hotspots is shown in bold.

| Method | PointVS | API |
| --- | --- | --- |
| Hotspots 1-5: $\Delta\text{SLE}_{20}$ | **0.304** | 0.213 |
| Hotspots 6-10: $\Delta\text{SLE}_{20}$ | 0.146 | **0.269** |

To test whether PointVS hotspots are also reliable for other proteins, we repeated the above process for two other targets: SARS-CoV-2 non-structural protein 3 (Mac1),^52^ and SARS-CoV-2 non-structural protein 14 (NSP14).^53^ These both have fragment screen datasets available on fragalysis^54,55^ (see SI Table 1). In both cases, these are considerably smaller than the Mpro dataset, and do not contain follow-up compounds.

Each of the top ten hotspots produced by PointVS and the API were tested, the results of which are shown in Table 7. Both sets of hotspots for Mac1 and NSP14 were tested on 35 and 30 fragments, respectively. The number of successfully elaborated fragments and mean number of elaborations generated per fragment for these targets are provided in SI Table 2.

**Table 7.**
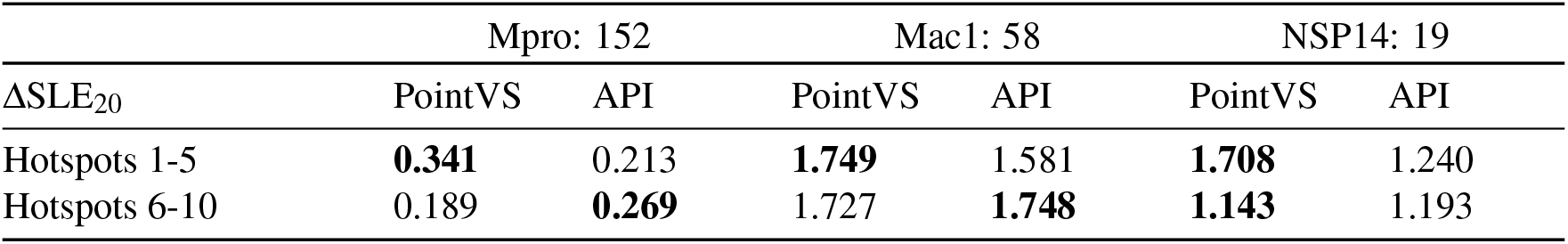
Mean standardised ligand efficiency of the top 20 elaborations (ΔSLE_20_) made using PointVS hotspots and API hotspots on three different targets: Mpro, Mac1, and NSP14. The numbers after the target names refer to the number of bound fragment structures available for each target. The list of fragalysis codes corresponding to the structures used is given in SI Table 1. The highest ΔSLE_20_ for both ranges of hotspots is shown in bold.

On average, the standardised ligand efficiency of molecules generated with PointVS hotspot maps is greater than those generated with the API hotspots. Within the PointVS scores, we see that the Mpro hotspots are lower scoring than those for the other two targets. This can be attributed to the 109 starting fragments for Mpro being of higher quality, having been extracted from curated follow-up compounds rather than fragments. The improvement that is made on them by elaborating is thus less significant. We also see that, unlike with the Hotspots API, the hotspots ranked 1-5 by PointVS consistently outperform those ranked 6-10, suggesting that the attribution method is good not only at picking out important atoms, but also ranking them amongst themselves.

## 4. Conclusion

In this paper we described the development of PointVS, an EGNN-based method for affinity prediction, and the first ML-based method for extracting important binding information from a target for molecule design.

During the development of PointVS we identified that a commonly used benchmark for MLBSFs, CASF-16, overestimates their accuracy when trained using the most commonly used training dataset. In light of this, PointVS was trained and tested on a rigorously filtered dataset, and hence encouraged to learn the rules governing intermolecular binding rather than memorise training data. We showed that when trained on this filtered dataset, PointVS achieves comparable results to other leading scoring functions, providing some evidence that it is successfully identifying important binding interactions. We provide further proof of this by performing attribution, and showing that PointVS is able to identify important interactions in line with those found by PLIP, a distance-based tool for profiling proteinligand interactions.

Finally, we investigated how knowledge of important binding regions could be leveraged in other stages of the drug development pipeline; namely fragment elaboration. We show that when provided with a set of bound structures, performing attribution on such structures yielded hotspot maps describing important protein atoms. Using these in unison with a fragment elaboration tool, STRIFE, resulted in improved docking scores for the elaborated molecules compared to when hotspots were obtained with a data-based method. This is further evidence that PointVS is not learning to memorise ligand information but to recognise important interactions, and additionally constitutes the first ML-based method of extracting structural information from a protein target in a way that is useful for fragment elaboration. More broadly, our work demonstrates that attribution techniques, when applied to debiased models, can be useful for extracting structural information in a way that can be useful in molecule generation.

## 5. Data and Software Availability

PointVS is available to download at https://github.com/oxpig/PointVS, as are our unbiased test and train splits.

## 6. Author Contributions

JS and CMD conceptualised and designed the study. All authors contributed to the methodology of the study. JS implemented the method. JS and LV conducted the experiments and analysed the results. NB, VC, PD, FD, and CMD supervised the project. All authors contributed to the writing, review, and editing of the manuscript.

## 7. Acknowledgements

JS was supported by funding from the Biotechnology and Biosciences Research Council (BB/S507611/1) and BenevolentAI, and LV was supported by funding from the Engineering and Physical Sciences Research Council (EP/W522211/1) and IBM Research. It was additionally supported by the Rosetrees Trust (Ref M940).

