## Supporting information for "A Step Towards Generalisability: Training a Machine Learning Scoring Function for Structure-Based Virtual Screening"

#### 1. Equivariance and Invariance

A function $\psi : X \to Y$ is *invariant* with respect to a group of transformations $G : X \to X$ if:

$$\psi(g(x)) = \psi(x) \quad \forall g \in G \quad \forall x \in X$$

$\psi$ is *equivariant* with respect to $G$ if there is a transformation $T : Y \to Y$ such that:

$$\psi(g(x)) = T(\psi(x)) \quad \forall g \in G \quad \forall x \in X$$

In other words, if $\psi$ is equivariant under $G$, applying elements of $G$ to $x$ will apply an analogous transformation to the output in $Y$. The order in which the transformations are applied does not matter to the end result.

While CNNs offer invariance to elements of the group $G = T(n)$, the set of all translations in $n$-dimensional space, they lack invariance to the group of all roto-translations, $SE(n)$. This means that rotating the protein-ligand input in 3D space may produce different predictions, behaviour which is undesirable. Usually the way to combat this is to augment training data by assigning random rotations to the inputs, but this does not achieve true rotational invariance. For a more thorough introduction to symmetry groups and equivariance, see references.[1–4]

Group-equivariant neural networks are a new class of deep learning model which offer invariance to various symmetry groups. They do this by using a series of equivariant layers followed by one or more invariant layer. For CNNs, these include Group Equivariant CNNs[1] and steerable CNNs.[5] CNNs still have the fundamental problem of a discrete input space and an extremely large input space and parameter count (a $48 \times 48 \times 48$ grid with 22 channels is more than $2.4 \times 10^6$ floating point numbers). If we view molecules as graphs, however, we can apply group-equivariant GNNs to the problem of pose predictions.

For these reasons, we use an architecture based on *E(n)*-Equivariant Graph Neural Network (EGNN) layers[2] for pose selection. The $E(n)$ symmetry group contains all elements from $SE(n)$, as well as the group of all reflections; EGNN layers are also permutation-equivariant, meaning that the network is invariant to the order of the input vertices.

### 2. PointVS: Methods

#### 2.1 Model

A graph is a natural way of representing a molecule, and attribution to individual inputs—namely atoms—is an intuitive process when using graph neural networks (GNNs). We use an architecture based on *E(n)*-Equivariant Graph Neural Network (EGNN) layers.[2] The $E(n)$ symmetry group contains all elements from $SE(n)$, as well as the group of all reflections; EGNN layers are also permutation-equivariant, meaning that the network is invariant to the order of the input vertices.

PointVS is a lightweight $E(n)$-equivariant graph neural network layer model, consisting of an initial projection to take the number of node features from 12 to 32, then 48 EGNN layers, followed by a global average pooling of the final node embeddings. This is followed by a sigmoid layer which gives a label $y \in [0,1]$ during the first stage of training on pose prediction. During finetuning on affinity data, this final layer is replaced by a randomly initiated fully connected layer and ReLU activation, which outputs $\mathbf{y} \in (\mathbb{R}^{+})^{3}$. This architecture includes residual connections for node features to allow for easier gradient flow and a richer combination of early and late representations. It also uses a shallow neural network as an attention mechanism,[6] which learns to score network edges—in this case, representing atomic interactions—by their importance.

#### 2.2 Input Features

Our EGNN takes four inputs: positions, node embeddings, edge indices, and edge embeddings. If there are $n$ atoms in an input structure, the positions tensor is an $n \times 3$ tensor containing the $x, y$ and $z$ coordinates of each atom. The node embeddings are an $n \times 12$ tensor of one-hot encoded OpenBabel atom types, with a bit to distinguish between ligand and receptor atoms.

Edges are generated on-the-fly using a variable distance cutoff. We use a cutoff of 10 Å for ligand-protein edges, and 2 Å for ligand-ligand and protein-protein edges. In this way, intramolecular edges closely mimic the covalent structure of input molecules, with a much more expansive connectivity to describe intermolecular interactions. The edge tensor is an $3 \times m$ matrix of $m$ edges, with one-hot encoding for ligand-ligand, ligand-receptor, and receptor-receptor interactions.

The binding pocket is defined as the collection of protein atoms within 6 Å of any ligand atom; the rest of the protein is ignored.

#### 2.3 Training

All models were trained for 10 epochs with a batch size of 32, using the Adam optimiser and cosine warm restarts with a maximum learning rate of $8 \times 10^{-4}$ and a weight decay of $10^{-4}$. For classification models, the selected model is the one with the best Top-1 value on the test set; for regression models, the selected model is the one with the highest Pearson's Correlation Coefficient (PCC) for affinity on the PDBBind Core Set.

For regression models with pretraining, once the classification task is complete, the best model is selected and the final linear layer with sigmoid is replaced by a multi-target regression, where the target is either $pK_d$, $pK_i$ or $pIC_{50}$. The labels for the two types of affinity data not present in the training set are ignored for calculating loss, so those weights are unaffected.

#### 2.4 Datasets

##### The Redocked Set

Training for pose classification uses the same training data as the Redocked model outlined by McNutt *et al.*[7] In previous work by the same authors,[8] ligands from the Pocketome v17.12[9] were redocked using AutoDock Vina into their cognate receptors, with an RMSD calculated between the redocked poses and the crystal pose. Sub-2 Å poses were labelled as active poses, with the rest labelled as inactive. Further negative examples were generated adversarially: a gnina CNN trained on the original redocked dataset was used to optimise docked poses. Optimised poses which were scored highly ($> 0.9$) by the trained CNN but were more than 2 Å RMSD from the crystal pose were added as inactive poses. In total, there are 52,385 active and 734,417 inactive poses in the version of Redocked2020 used here, which we have shortened to Redocked.

##### PDBBind Refined: gninaSet$_{Pose}$ Set

The PDBBind refined set v.2019[10] is a carefully selected set of high-quality protein-ligand complexes, complete with binding affinities. In their gnina 1.0 paper,[8] the authors took the 4,852 unique protein-ligand complexes, removing solvents and any other molecules besides the protein and the ligand, as well as removing structures with ligands that do not have a mass between 150 and 1,000 Da. This filtered, preprocessed version of the refined set is hosted by the authors of gnina 1.0[8] at `https://bits.csb.pitt.edu/files/gnina1.0_paper`. Gnina was used with default settings, excepting `--num_modes=50`, `--min_rmsd_filter=0.0`, and the use of the standard Vina scoring function in refinement and ranking rather than a CNN.

Of the 4,260 filtered structures, the 441 which are not used in training the 'redocked' gnina model are selected as the test set for consistency with that work, which we call gninaSet$_{Pose}$. Up to 50 poses are reranked by our GNN (some of which may be very similar), after which the standard 1.0 Å RMSD filter is applied to get a final pose ranking.

**PDBBind General and Core Sets**

The PDBBind v.2020 General set contains protein-ligand complexes and binding affinities curated from the PDB.[11] The affinities are in the form of $pK_d$, $pK_i$ or $pIC_{50}$, and this was the training set for affinity regression. The PDBBind Core set, also known as the CASF-2016 test set for affinity regression, comprises 285 high-quality crystal structures, as well as their binding affinities,[12] and is a subset of the General set. This is the test set for regression (*scoring power*).

#### 3. Assessing Attribution Methods

As described in the paper, we use three methods for attribution: atom masking, bond masking, and edge attention analysis. We showed that when carrying out atom masking with gnina,[8] bond masking with InteractionGraphNet (IGN)[13] and edge attention analysis with PointVS, the method that extracted important protein atoms in most agreement with PLIP, the protein-ligand interaction profiler.[14] To learn to what extent this could be accredited to the method used for attribution, we also carried out atom masking and bond masking on PointVS, the results of which are given in Table 1.

| Attribution | MLBSF | $\rho_5$ | $\rho_{10}$ |
|---|---|---|---|
| Edge attention | PointVS | 0.640 | 0.788 |
| Atom masking | gnina | 0.234 | 0.093 |
| | PointVS | 0.152 | 0.169 |
| Bond masking | IGN | 0.209 | -0.071 |
| | PointVS | 0.140 | 0.122 |

Table 1. Mean rank correlation calculated between the scores of the top five ($\rho_5$) and top ten ($\rho_{10}$) ranked protein atoms by PointVS, gnina, and InteractionGraphNet (IGN) and the distance between them and the nearest polar ligand atom. The means were calculated over a subset of 20 randomly selected bound structures from the PDBBind Core set (see SI for PDB IDs).

#### 4. Hotspots API: Calculation and Processing of Hotspots

For a given protein target, the API calculates acceptor, donor and apolar atomic propensity maps using SuperStar.[15] This tool defines a grid covering the protein with points spaced 0.5Å apart, and then using data from the Cambridge Structural Database (CSD),[16] assigns a propensity for the three atomic probe types—acceptor, donor, and apolar—to each point. These scores are assigned based on the principle that if a positive interaction between two groups at a certain distance and angle is made, then it will occur more frequently in structures stored in the CSD, and thus be assigned a higher propensity score. These grid point scores are then weighted according to how buried in the protein they are, so that more buried sites are favored.

Further scores are then calculated with small chemical probes consisting of aromatic rings with an acceptor, donor, or apolar atom in the substituent position. The three weighted propensity grids are sampled by their corresponding probe: a given probe is translated to all grid points with scores above a threshold of 15, and randomly rotated 3000 times around the centre of the substituent atom. For each pose, the probe atom scores are read from their corresponding grids, and the probe score is found as the geometric mean of the scores of the probe's atoms. Once complete, the sampled probe scores are assigned to an output grid.

To process the donor and acceptor grids, we follow the protocol outlined in Hadfield *et al.*[17] A greedy clustering algorithm is employed. A cluster is initialised as a random single point, and all points less than 1Å away that are not part of another cluster are added. Then, for each point added, this process is repeated, so all remaining unclustered points within 1Å are added. This process continues until there are no unclustered points within 1Å of the points forming the cluster. Once a cluster is finalised, a new cluster is defined by selecting another single unclustered point. This process is repeated until all points have been assigned to a cluster. The centroids of these clusters are defined as the mean position of the points constituting them. Clusters comprising less than eight points are removed, as well as those with centroids outside of an apolar region (this is assessed using the apolar grid). The remaining clusters are then allocated a score consisting of the number of points comprising them, and output alongside their associated scores as the acceptor and donor hotspot maps for a given protein target.

| Mpro | | | Mac1 | | NSP14 |
|---|---|---|---|---|---|
| x10474_0A | x10484_0A | x10494_0A | DLS-EU0569_0A | DLS-X0626_0A | x1203_0D |
| x10898_0A | P0906_0B | x11543_0A | DLS-X0724_0A | DLS-X0689_0A | x1088_0D |
| x2964_0A | x11426_0A | x2646_0A | DLS-X0591_0A | DLS-X0711_0A | x1201_0D |
| x11372_0A | x10604_0A | x10559_0A | DLS-X0681_0A | DLS-EU0034_0A | x1316_0D |
| x11233_0A | x12682_0A | x11764_0A | DLS-X0895_1A | DLS-EU0087_0A | x1448_0D |
| x10473_0A | x10889_0A | x2971_0A | DLS-X0910_0A | | x1412_0D |
| x10327_0A | P1470_0A | x11159_0A | DLS-X0587_0A | | x1181_0D |
| x12740_0A | x10723_0A | x2908_0A | DLS-X0421_0A | | x1199_0D |
| x10888_0A | x10248_0A | x11532_0A | DLS-EU0342_0A | | x1197_0D |
| x10371_0A | P0045_0B | x0678_0A | DLS-X0844_0A | | x1384_1D |
| x10606_0A | x10756_0A | x11164_0A | DLS-EU0815_0A | | x1486_0D |
| x11743_0A | x10995_0A | P0045_0A | DLS-EU0291_0A | | x1245_0D |
| x2912_0A | x10555_0A | x3108_0A | DLS-X0299_0A | | x1541_0D |
| x10565_0A | x2572_0A | x11339_0A | DLS-X0471_0A | | x1603_0D |
| x10422_0A | P0906_0A | x11564_0A | DLS-X0895_0A | | x1236_2D |
| x10421_0A | P2007_0B | x10789_0A | DLS-EU0238_0A | | x1282_0D |
| P1073_0A | x3359_0A | P2001_0B | DLS-X0766_0A | | x1641_0D |
| x10488_0A | x10387_0A | x2608_0A | DLS-X0926_0A | | x1078_0D |
| x11427_0A | x0107_0A | x10856_0A | DLS-EU0056_0A | | x1212_1D |
| P0145_0B | x10201_0A | x10314_0A | DLS-X0592_0A | | |
| x10942_0A | x11271_0A | x11513_0A | DLS-X0967_0A | | |
| x10022_0A | x2581_0A | x11540_0A | DLS-EU0144_0A | | |
| x11473_0A | x10377_0A | x2600_0A | DLS-X0722_0A | | |
| x0434_0A | x10395_0A | x10598_0A | DLS-EU0704_0A | | |
| P1470_0B | x10329_0A | x11041_0A | DLS-EU0844_0A | | |
| x12719_0A | x10178_0A | x11417_0A | DLS-EU0481_0A | | |
| x10396_0A | x12300_0A | x10638_0A | DLS-X0969_0A | | |
| z7vh8_0A | x10324_0A | x10334_0A | DLS-EU0241_0A | | |
| x11368_0A | P0884_0A | x10392_0A | DLS-X0962_0B | | |
| P2007_0A | x11346_0A | x11708_0A | DLS-X0742_0A | | |
| x11231_0A | x2649_0A | x11318_0A | DLS-EU0172_0A | | |
| x10906_0A | x11641_0A | x10019_0A | DLS-X0918_0A | | |
| x3080_0A | x10800_0A | x11428_0A | DLS-X1018_0A | | |
| x11560_0A | x10996_0A | x10733_0A | DLS-X0962_0A | | |
| x11013_0A | x10417_0A | x11475_0A | DLS-X0900_0A | | |
| x10566_0A | x10247_0A | x10478_0A | DLS-X0685_0A | | |
| P2001_0A | x3298_0A | x11485_0A | DLS-X0962_1A | | |
| x11809_0A | x3366_0A | x10575_0A | DLS-X0805_0A | | |
| x10679_0A | x2643_0A | P0145_0A | DLS-X0142_0A | | |
| x10801_0A | x11488_0A | x10787_0A | DLS-X0104_0A | | |
| P0925_0A | x11317_0A | x2569_0A | DLS-X0852_0B | | |
| x11225_0A | x10889_1A | x11044_0A | DLS-X0918_1A | | |
| x10338_0A | x10476_0A | x11723_0A | DLS-EU0716_0A | | |
| x10359_0A | x11562_0A | x10322_0A | DLS-EU0123_0A | | |
| x2193_0A | x10900_0A | x11354_0A | DLS-X0905_0A | | |
| x10423_0A | x2562_0A | x12321_0A | DLS-X0910_1A | | |
| x11557_0A | x11801_0A | x11025_0A | DLS-X0933_0A | | |
| x10645_0A | x10237_0A | x10834_0A | DLS-X0837_0A | | |
| x10626_0A | x11011_0A | x11507_0A | DLS-EU0293_0A | | |
| x10236_0A | x10535_0A | x10976_0A | DLS-EU0424_0A | | |
| x11186_0A | x10610_0A | | DLS-EU0046_0A | | |

Table 2. Fragalysis identification codes of the structures supplied to PointVS to obtain hotspots.

#### 5. Structure IDs

The bound fragment structures used for all three targets (Mpro, Mac1 and NSP14) are listed in Table 2. These datasets are all available to download from the fragalysis platform.[18–21]

The PDB IDs of the randomly selected subset of 20 structures from the PDBBind core set used in section 3.4.2 are 3URI, 4DE1, 4DLI, 3N76, 4M0Z, 3TWP, 1VSO, 3NQ9, 3UEU, 3AO4, 3JVR, 2BRB, 1H23, 1E66, 2FVD, 2ZB1, 4K18, 3OE5, 3GE7, 3RR4, 3UI7, 4JSZ, 3QQS, 2VKM, 4MGD, 2XII, 2QE4, 1Y6R.

#### 6. Mpro: Hotspot Map Dependency on Fragment Screen Size

We randomly generate sets of bound structures of Mpro of various sizes, and perform attribution on them to obtain PointVS hotspots. The results from this are shown in Fig 4.

As another assessment of the consistency of PointVS hotspots, we assert that the top five hotspots found when PointVS is given all of the Mpro fragment screen structures in Table 1 can be considered the 'ground truth' top five hotspots. We then carry out attribution on various sizes of randomly sampled sets of bound structures, as above, and record how many of the top five hotspots for a given random sample coincide with the ground truth top five hotspots. The results from this are shown in Fig 1. We see that increasing the size of the randomly selected set of bound structures results in increased agreement with the ground truth hotspots. However, we also note that this effect drops off when the size of the sets of bound structures reaches 30, implying that after this point, increasing the size of a fragment screen would have little benefit on the quality of the top five hotspots produced by PointVS. This is a promising sign of the applicability of the PointVS hotspots, as Mpro is very unusual in having such a large number of fragment structures available.

#### 7. Fragment Elaboration: Numbers of Elaborations Generated

Hotspots for the targets Mac1 and NSP14 are tested on 35 and 30 fragments respectively. The number of fragments successfully elaborated for each hotspot and the mean number of elaborations for each fragment are shown in Tables 3a and b.

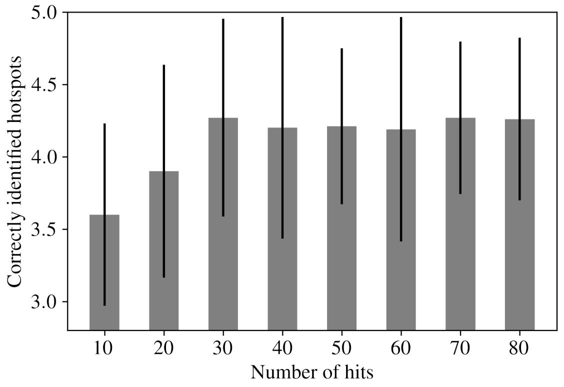


Fig. 1. The average number of hotspots found by PointVS using different numbers of fragment structures of Mpro that agree with those found using the full set of 152 fragment structures, with error bars showing the standard deviation of the number of ground truth hotspots identified.

#### 8. FGFR1 Case Study

To demonstrate the potential of the PointVS hotspot maps for real-world fragment-to-lead campaigns, we present a case study on a well-studied set of drug targets: the fibroglast growth factor receptors. Aberrant activation of these receptors is associated with a wide variety of cancers, making them a promising druggable target. In 2019, a collaboration between Astex and Newcastle University resulted in erdafitnib:[22] a drug that has since been FDA approved, making it the third fragment-derived drug to reach this stage.

In the process of designing erdafitnib, the collaborators heavily relied on a fragment screen performed on the target in 2008. A fragment found in this screen, shown in Fig 2, was found to have high potency, and was used as a starting point in the design process.

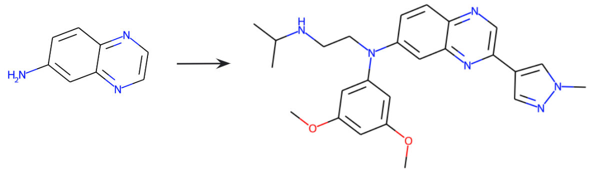

Fig. 2. Left, the fragment used as a starting point in the development of erdafitnib, shown on the right.

In Patani *et al.*,[23] two hydrogen bonds are found to form between erdafitnib and the protein. We investigated whether we could identify the protein atoms forming these bonds as important using either PointVS or the API hotspots.

The API placed one donor hotspot 1.5Å away from the location of one of the erdafitnib atoms that forms a hydrogen bond with a protein atom. However, it did not recognise the point the second binding ligand atom is found at as important.

To obtain the PointVS hotspots, we performed attribution on 18 of the bound structures of FHFR1 available in the PDB.[11] These bound structures contain compound-like molecules rather than fragments, so we filtered them to ensure that none had a Tanimoto similarity of greater than 0.7 to erdafitnib. This resulted in us selecting the structures corresponding to the following PDB codes: 4NKS, 4NK9, 1AGW, 3C4F, 4NKA, 4V01, 4V05, 5B7V, 5A46, 4UWB, 3RHX, 1FGI, 5A4C, 4UWC, 5VND, 6P69.

We found that PointVS correctly identified both of the protein atoms that form hydrogen bonds with ligand atoms as important, ranking them as the third and sixth most important atoms for binding.

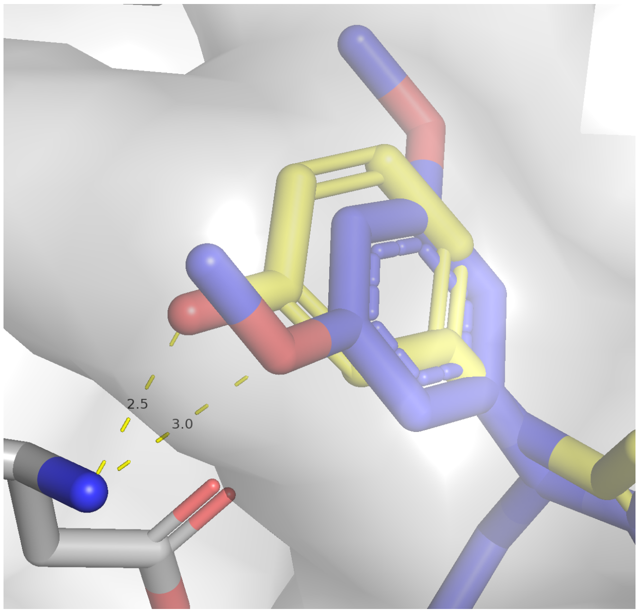


Fig. 3. Elaboration generated by STRIFE using the PointVS hotspots (yellow), that recovers the same hydrogen bond as erdafitnib, shown in blue (PDB ID: 5EW8[23]).

To provide an example of how knowledge of these two hotspots could be used, we supplied one of them to STRIFE to demonstrate that molecules similar to erdafitnib could be generated. An example of an elaboration that recovered the same binding mode as erdafitnib is shown in Fig 3.

[11] S. K. Burley, C. Bhikadiya, C. Bi, S. Bittrich, L. Chen, G. V. Crichlow, C. H. Christie, K. Dalenberg, L. Di Costanzo, J. M. Duarte, S. Dutta, Z. Feng, S. Ganesan, D. S. Goodsell, S. Ghosh, R. K. Green, V. Guranović, D. Guzenko, B. P. Hudson, C. L. Lawson, Y. Liang, R. Lowe, H. Namkoong, E. Peisach, I. Persikova, C. Randle, A. Rose, Y. Rose, A. Sali, J. Segura,

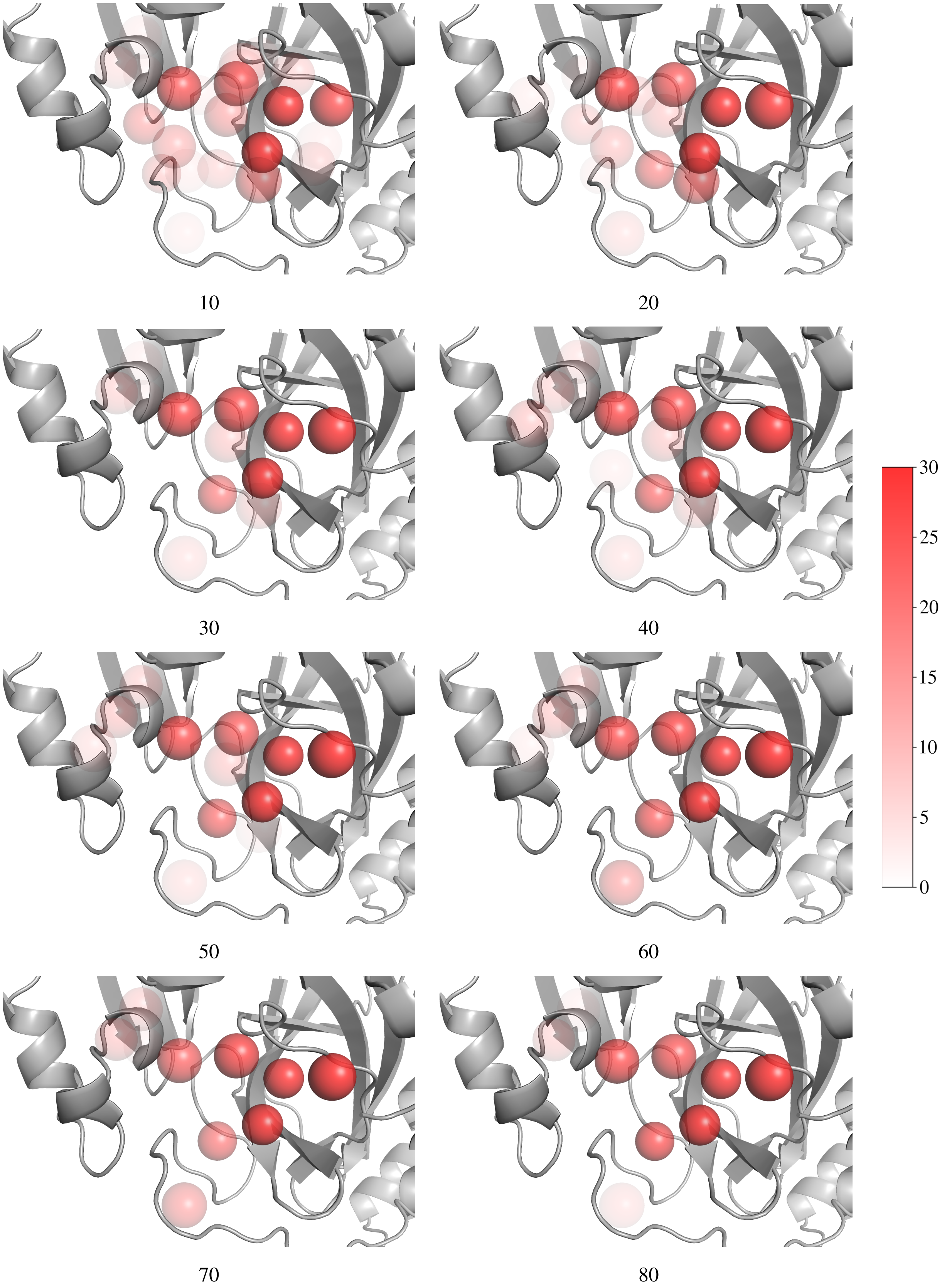


Fig. 4. Top 5 PointVS donor and acceptor hotspots (both shown in red) obtained in the pocket of Mpro by performing attribution on 40 randomly generated sets of bound fragments. The number of structures constituting each random set is shown below each image. Darker spheres correspond to hotspots which were included in the top 5 more frequently.

| | PointVS | | Hotspots API | |
|---|---|---|---|---|
| Rank | Fragments successfully elaborated | Elaborations per fragment | Fragments successfully elaborated | Elaborations per fragment |
| 1 | 21 | 217 | 15 | 189 |
| 2 | 12 | 187 | 10 | 183 |
| 3 | 13 | 223 | 11 | 207 |
| 4 | 11 | 192 | 7 | 158 |
| 5 | 5 | 207 | 12 | 179 |
| 6 | 9 | 176 | 15 | 162 |
| 7 | 6 | 239 | 21 | 220 |
| 8 | 2 | 222 | 11 | 197 |
| 9 | 0 | 0 | 0 | 0 |
| 10 | 1 | 218 | 8 | 210 |

a

| | PointVS | | Hotspots API | |
|---|---|---|---|---|
| Rank | Fragments successfully elaborated | Elaborations per fragment | Fragments successfully elaborated | Elaborations per fragment |
| 1 | 14 | 194 | 2 | 237 |
| 2 | 3 | 196 | 23 | 218 |
| 3 | 24 | 215 | 24 | 195 |
| 4 | 23 | 186 | 14 | 154 |
| 5 | 14 | 183 | 0 | 0 |
| 6 | 23 | 186 | 24 | 195 |
| 7 | 17 | 194 | 0 | 0 |
| 8 | 5 | 196 | 4 | 116 |
| 9 | 1 | 89 | 27 | 199 |
| 10 | 6 | 205 | 13 | 181 |

b

Table 3. The number of fragments that were successfully elaborated on using PointVS and API hotspots for Mac1 (a) and NSP14 (b), and the mean number of elaborations generated per fragment.

M. Sekharan, C. Shao, Y.-P. Tao, M. Voigt, J. D. Westbrook, J. Y. Young, C. Zardecki, and M. Zhuravleva, "RCSB Protein Data Bank: powerful new tools for exploring 3D structures of biological macromolecules for basic and applied research and education in fundamental biology, biomedicine, biotechnology, bioengineering and energy sciences," *Nucleic Acids Research*, vol. 49, pp. D437–D451, 11 2020.
